# Calreticulin co-expression supports high level production of a recombinant SARS-CoV-2 spike mimetic in *Nicotiana benthamiana*

**DOI:** 10.1101/2020.06.14.150458

**Authors:** Emmanuel Margolin, Matthew Verbeek, Ann Meyers, Ros Chapman, Anna-Lise Williamson, Edward P Rybicki

## Abstract

An effective prophylactic vaccine is urgently needed to protect against SARS-CoV-2 infection. The viral spike, which mediates entry into cells by interacting with the host angiotensin-converting enzyme 2, is the primary target of most vaccines in development. These vaccines aim to elicit protective immunity against the glycoprotein by use of inactivated virus, vector-mediated delivery of the antigen *in vivo*, or by direct immunization with the purified antigen following expression in a heterologous system. These approaches are mostly dependent on the growth of mammalian or insect cells, which requires a sophisticated infrastructure that is not generally available in developing countries due to the incumbent costs which are prohibitive. Plant-based subunit vaccine production has long been considered as a cheaper alternative, although low expression yields and differences along the secretory pathway to mammalian cells have posed a challenge to producing certain viral glycoproteins. Recent advances that have enabled many of these constraints to be addressed include expressing the requisite human proteins in plants to support the maturation of the protein of interest. In this study we investigated these approaches to support the production of a soluble and putatively trimeric SARS-CoV-2 spike mimetic in *Nicotiana benthamiana* via transient *Agrobacterium-mediated* expression. The co-expression of human calreticulin dramatically improved the accumulation of the viral spike, which was barely detectable in the absence of the co-expressed accessory protein. The viral antigen was efficiently processed even in the absence of co-expressed furin, suggesting that processing may have occurred at the secondary cleavage site and was mediated by an endogenous plant protease. In contrast, the spike was not efficiently processed when expressed in mammalian cells as a control, although the co-expression of furin improved processing considerably. This study demonstrates the feasibility of molecular engineering to improve the production of viral glycoproteins in plants, and supports plant-based production of SARS-CoV-2 spike-based vaccines and reagents for serological assays.

## Introduction

The absence of suitable infrastructure to produce vaccines or diagnostics to combat emerging viruses leaves most developing countries vulnerable and unable to respond appropriately to emerging disease threats. This was highlighted during the 2009 H1N1 pandemic, and remains a major challenge to this day, as evidenced by the ongoing SARS-CoV-2 pandemic [1]. Whilst there are currently over 100 vaccines in preclinical development and 10 in clinical testing, none of these are being developed in Africa, and concerns have been raised as to whether the current world manufacturing capacity will be sufficient to accommodate the global demand for a SARS-CoV-2 vaccine [2].

Following the 2009 H1N1 pandemic, it was acknowledged that the accepted vaccine manufacturing paradigm was not equipped to contend with pandemic outbreaks due to limitations of production scale and slow development time lines [2]. Plant-based vaccine production was therefore proposed as an alternative platform for vaccine development, due to the potential for rapid large scale antigen production [3]. The system is also well-suited to developing countries where financial resources are often lacking, as the costs to establish a manufacturing facility and to produce pharmaceuticals are lower than those required for conventional mammalian cell production platforms [4]. Accordingly, several candidate influenza subunit vaccines have been produced in plants and have subsequently been tested in clinical trials [5–7]. These vaccines are based on the viral haemagglutinin glycoprotein, which accumulates at high levels in plants and elicits protective immunity, despite not being efficiently proteolytically cleaved in the system [8]. A quadrivalent plant-produced influenza vaccine produced by Medicago Inc. is currently under consideration by Health Canada [9].

Many other viral glycoproteins, however, do not accumulate at such high levels in plants and the host cellular machinery may not adequately support critical folding and processing events that are required for glycoprotein maturation [3]. Furin, the enzyme responsible for the proteolytic processing of many viral glycoproteins, is not naturally produced in plants [10]. Additional constraints, such as the host chaperone machinery or differences in the plant oligosaccaryltransferase complex, may also complicate the production of, and glycosylation of, certain viral glycoproteins *in planta* [11–13]. Recently, it has been proposed that engineering the plant secretory pathway to support the production of viral glycoproteins could improve their accumulation in plants, and support their maturation during synthesis [9]. The co-expression of human chaperones has recently been reported to improve the accumulation of a range of envelope viral glycoproteins in *N. benthamiana*, including the HIV envelope glycoprotein and the Epstein-Barr gp350 glycoprotein, which are possibly amongst the most complex proteins produced in a plant expression system to date [13]. This approach was also combined with the expression of human furin to support proteolytic processing *in planta*, where this would not otherwise occur [13].

Several plant biotechnology companies have released press statements reporting the production of candidate SARS-CoV-2 vaccines in *N. benthamiana*, although it is not clear which antigens are being produced or which expression approach is being employed. The viral spike is the logical target for vaccine development as the glycoprotein is critical for the infection of target cells [14]. The spike comprises of a heavily glycosylated trimer that initiates infection by engaging the host angiotensin converting enzyme 2 [15, 16]. In addition to glycosylation, the spike also undergoes proteolytic maturation by furin proteases during its synthesis [17]. It is presently unclear if the spike glycoprotein can be produced at high levels in plants, if it requires further modification to accumulate at reasonable levels, or if it will be appropriately folded and processed along the secretory pathway. Therefore, in this study we explored the co-expression of human chaperones and the protease furin to support the production of a soluble SARS-CoV-2 spike glycoprotein, with the intention of developing capacity to cheaply produce vaccine antigens and diagnostic reagents for further development.

## Results

### Modification of the SARS-CoV-2 spike sequence for heterologous expression

A soluble spike antigen (SΔTM), consisting of both the S1 and S2 subunits but lacking the transmembrane and cytoplasmic domains, was designed for optimal expression and processing in plants (**Figure 1A**). The modified spike included an optimized cleavage sequence (RRRRRR) to promote processing by furin, as previously reported for the HIV envelope glycoprotein when produced in both plants and mammalian cells [13, 18]. The synthesized sequence was also human codon optimized for dual expression in human and plant cells, resulting in a 56% GC content. This codon optimization strategy was informed by several other studies where human codon optimization yielded higher expression than the corresponding plant codon usage [1, 19, 20].

**Figure 1:**
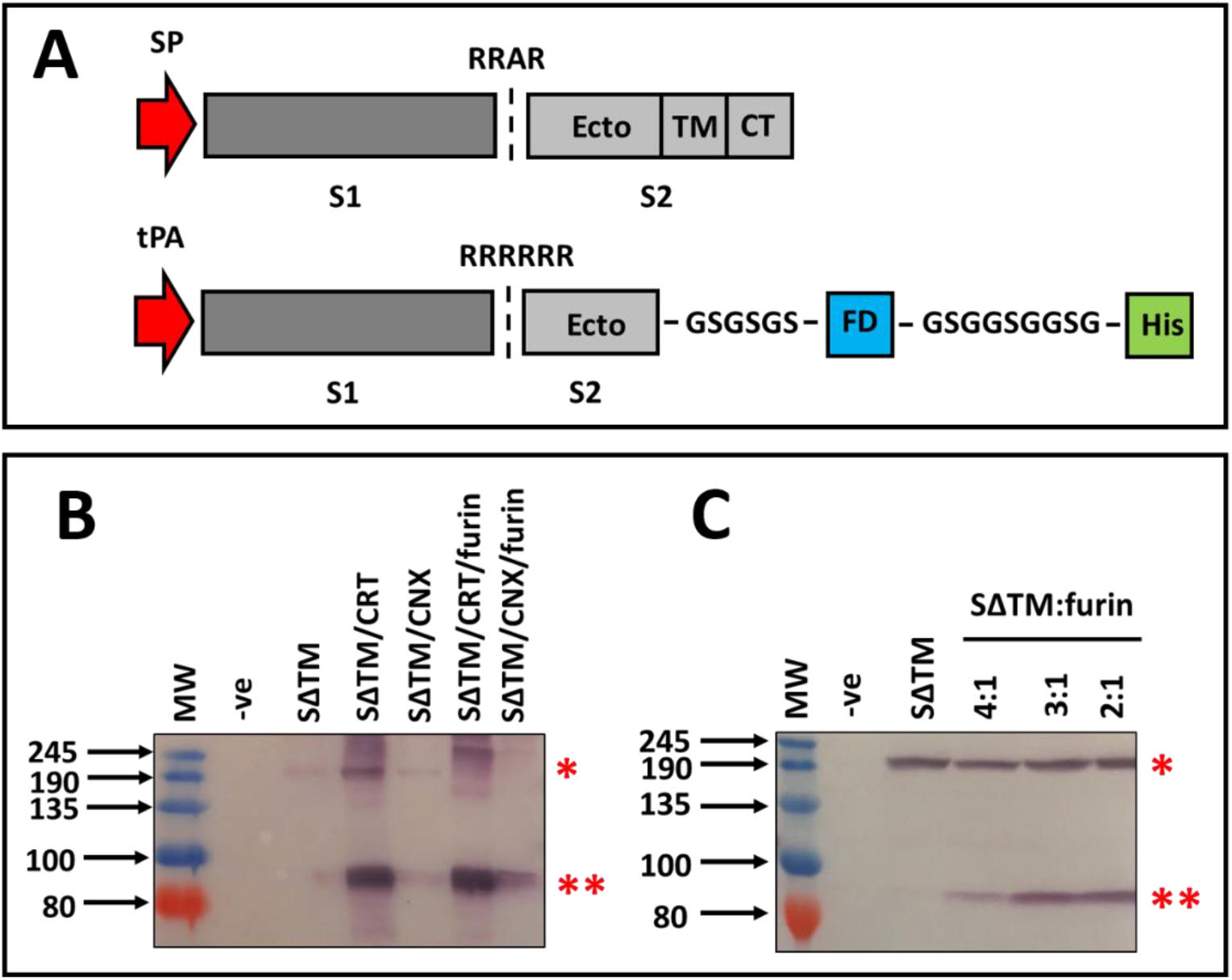
Design and expression of an engineered SARS-CoV-2 spike mimetic. A) Schematic of the native spike glycoprotein (top) and modified SΔTM antigen (bottom) that was designed for heterologous expression. The S1 and S2 subunits of the spike are denoted as dark and light grey boxes respectively. The leader sequence is indicated by a red arrow. The furin cleavage site is indicated by the vertical dotted line the sequence indicated above. The GCN4 foldon (FD) is indicated as a blue box. The polyhistidine tag (His) is shown as a green box. (SP=native signal peptide, tPA= tissue plasminogen activator leader sequence, Ecto = ectodomain of S2, TM= transmembrane domain). B) Western blotting of plant lysate to detect expression of the SARS-CoV-2 spike antigen following agroinfiltration. A stroke (/) indicates where more than 1 protein was co-expressed. (-ve=uninfiltrated, SΔTM=soluble spike antigen, CRT=calreticulin, CNX=calnexin). C) Western blotting of transfected mammalian lysate following transient expression of the recombinant spike protein. The recombinant spike protein was detected using polyclonal mouse anti-His antibody. In both B) and C) the uncleaved product is indicated by a * and the cleaved product is indicated by a **.

### Expression of the SARS-CoV-2 spike in *Nicotiana benthamiana*

Three different transient expression strategies were investigated to produce the SARS-CoV-2 spike in plants. These included the transient expression of the protein alone, co-expression with each of the lectin binding chaperones calreticulin (CRT) and calnexin (CNX), and the co-expression of the spike with each of the chaperones and human furin. Recombinant *A. tumefaciens* encoding the heterologous proteins were vacuum infiltrated into *N. benthamiana* and leaf-derived protein extract was subjected to western blotting to confirm expression of the recombinant protein (**Figure 1B**). In the absence of any co-expressed chaperone, a faint band corresponding to the uncleaved spike (SΔTM) protein was detected above the 190 kDA marker. In contrast, calreticulin co-expression resulted in a clear product at this size, as well as a band between 80 kDa and 100 kDA which corresponds to the expected S2 cleavage product. Processing occurred in the absence of co-expressed furin and was not improved when the protease was co-expressed with the glycoprotein and the chaperone. Calnexin co-expression did not appear to improve the accumulation of the spike.

### Transient expression of the SARS-CoV-2 spike in mammalian cells

The SΔTM antigen was also expressed in mammalian cells as a control to verify that the products from *N. benthamiana* were of comparable size. This also served as a proxy to determine if there was evidence of under glycosylation of the protein in plants, as has been described for other proteins [12]. In addition, the expression plasmid encoding the spike was also co-transfected with a second plasmid encoding furin at increasing ratios. Western blotting of lysate from transfected cells confirmed the expression of the recombinant protein and yielded products of the same size as those observed in plants (**Figure 1C**). In contradiction to other published results, the protein was predominantly unprocessed in mammalian cells despite including a modified site for improved processing [17]. Co-expression of furin did however improve processing, yielding the expected cleavage product, which increased in intensity with higher levels of furin expression.

## Discussion and conclusions

This study describes the first publicly available account detailing the expression of the SARS-CoV-2 spike in a plant-based expression system. Consistent with other studies reporting inefficient production of viral glycoproteins in plants, we noted that the spike glycoprotein accumulates at low levels in plants using routine expression approaches [3]. However, the co-expression of human CRT, as was also the case with soluble HIV Env trimers [13], improved the accumulation of the protein considerably, enabling the production of the antigen at sufficient levels to warrant further development as a vaccine antigen or as a diagnostic reagent. Whilst the soluble protein benefitted from co-expression of CRT, production of the full-length spike may require the co-expression of CNX which preferentially mediates the folding of membrane-bound glycoproteins. Other comparable soluble spike antigens are also in development, such as the stabilized prefusion trimer developed by the University of Queensland. Interestingly, the spike was efficiently processed in plants, despite the lack of furin production along the secretory pathway, and processing was not improved following the co-expression of the protease. This may be due to recognition of a secondary cleavage site by an endogenous plant protease. The sizes of the recombinant protein products produced in mammalian cells and plants were indistinguishable following western blotting, and did not suggest that under glycosylation of the antigen would be a concern. However, further experiments are required to confirm this. This work lends credence to developing molecular engineering approaches to support the production of complex biologics in plants, and may have similar value to support the production of other SARS-CoV-2 spike glycoprotein-based antigens and diagnostics that are being pursued in plant systems.

## Experimental procedures

### Antigen design and generation of recombinant expression plasmids

A soluble derivative of the SARS-CoV-2 spike was designed based on the first publicly available sequence (GenBank accession: MN908947.3). The coding sequence was truncated to remove the transmembrane and cytoplasmic domains of the glycoprotein. The native leader sequence was replaced with the tissue plasminogen activator (tPA) signal peptide and the putative furin recognition sequence (RRAR) was replaced with a hexa-arginine (RRRRRR) motif to promote proteolytic processing. A GCN4 Fibritin trimerization motif was incorporated at the end of the gene sequence preceded by a flexible linker peptide (GSGSGS). A polyhistidine (HHHHHH) affinity tag was added to the C-terminal end of the antigen after a second linker (GSGGSGGSG). The antigen coding sequence was synthesized by GenScript to reflect the preferred human codon usage and synthetic restriction sites were added to the 5’ (HindIII and AgeI) and 3’ (XhoI and EcoRI) termini of the DNA. The gene sequence was cloned into pEAQ-*HT*, using AgeI and XhoI, for expression in plants [21]. The recombinant plasmid was then transformed into *A tumefaciens* AGL1 as previously described [13]. Recombinant *A. tumefaciens* encoding human chaperones and furin were previously described [13]. The spike antigen sequence was also cloned into pTHpCapR, using HindIII and EcoRI, for expression in mammalian cells [22]. The furin expression construct used in this study was previously described [13].

### Recombinant protein production in mammalian cells

Protein expression as conducted in adherent HEK293 cells. Cells were maintained in DMEM (1×) + GlutaMAX^™^-1 (Gibco) supplemented with 10% fetal bovine serum (Gibco) and Penicillin/Streptomycin (Biowhittaker^®^, Lonza). Transfections were conducted using X-tremeGENE^™^ HP DNA Transfection Reagent in accordance with the manufacturer’s instructions. Crude protein lysate was harvested 72 hours post transfection using Glo Lysis Buffer.

### Recombinant protein production in *Nicotiana benthamiana*

Protein expression in plants was conducted as previously described [13]. In the case where multiple proteins were co-expressed, equal amounts (OD600) of the relevant bacterial cultures were mixed. Crude plant homogenate was harvested 3 days post-infiltration, using Tris-buffered saline [pH 8.0], as previously detailed [19].

### Immunoblotting to detect expression of the SARS-CoV-2 spike

Equal volumes of total soluble protein were resolved on SDS-PAGE gels and then immunoblotted using established procedures [13, 19]. The recombinant spike was detected with 1:2000 of mouse monoclonal anti-histidine antibody (Serotech, MCA1396), which in turn was detected using 1:5000 goat anti-mouse IgG alkaline phosphatase conjugate (Sigma, A3562).

## References

1. Mortimer, E., et al., Setting up a platform for plant-based influenza virus vaccine production in South Africa. BMC Biotechnol, 2012. 12: p. 14.

2. Mullard, A., COVID-19 vaccine development pipeline gears up. Lancet, 2020. 395(10239): p. 1751–1752.

3. Margolin, E., et al., Production of complex viral glycoproteins in plants as vaccine immunogens. Plant Biotechnol J, 2018.

4. Murad, S., et al., Molecular Pharming for low and middle income countries. Curr Opin Biotechnol, 2019. 61: p. 53–59.

5. Cummings, J.F., et al., Safety and immunogenicity of a plant-produced recombinant monomer hemagglutinin-based influenza vaccine derived from influenza A (H1N1)pdm09 virus: a Phase 1 dose-escalation study in healthy adults. Vaccine, 2014. 32(19): p. 2251–9.

6. Pillet, S., et al., A plant-derived quadrivalent virus like particle influenza vaccine induces cross-reactive antibody and T cell response in healthy adults. Clin Immunol, 2016. 168: p. 72–87.

7. Pillet, S., et al., Immunogenicity and safety of a quadrivalent plant-derived virus like particle influenza vaccine candidate-Two randomized Phase II clinical trials in 18 to 49 and >/=50 years old adults. PLoS One, 2019. 14(6): p. e0216533.

8. D’Aoust, M.A., et al., Influenza virus-like particles produced by transient expression in Nicotiana benthamiana induce a protective immune response against a lethal viral challenge in mice. Plant Biotechnol J, 2008. 6(9): p. 930–40.

9. Margolin, E., et al., Engineering the Plant Secretory Pathway for the Production of Next-Generation Pharmaceuticals. Trends Biotechnol, 2020.

10. Wilbers, R.H., et al., Co-expression of the protease furin in Nicotiana benthamiana leads to efficient processing of latent transforming growth factor-beta1 into a biologically active protein. Plant Biotechnol J, 2016. 14(8): p. 1695–704.

11. Göritzer, K., et al., Efficient N-Glycosylation of the Heavy Chain Tailpiece Promotes the Formation of Plant-Produced Dimeric IgA. Frontiers in Chemistry, 2020. 8(346).

12. Castilho, A., et al., An oligosaccharyltransferase from Leishmania major increases the N-glycan occupancy on recombinant glycoproteins produced in Nicotiana benthamiana. Plant Biotechnol J, 2018. 16(10): p. 1700–1709.

13. Margolin, E., et al., Co-expression of human calreticulin significantly improves the production of HIV gp140 and other viral glycoproteins in plants. Plant Biotechnol J, 2020.

14. Amanat, F. and F. Krammer, SARS-CoV-2 Vaccines: Status Report. Immunity, 2020.

15. Hoffmann, M., et al., SARS-CoV-2 Cell Entry Depends on ACE2 and TMPRSS2 and Is Blocked by a Clinically Proven Protease Inhibitor. Cell, 2020.

16. Song, W., et al., Cryo-EM structure of the SARS coronavirus spike glycoprotein in complex with its host cell receptor ACE2. PLoS Pathog, 2018. 14(8): p. e1007236.

17. Walls, A.C., et al., Structure, Function, and Antigenicity of the SARS-CoV-2 Spike Glycoprotein. Cell, 2020.

18. Binley, J.M., et al., Enhancing the proteolytic maturation of human immunodeficiency virus type 1 envelope glycoproteins. J Virol, 2002. 76(6): p. 2606–16.

19. Margolin, E., et al., Production and Immunogenicity of Soluble Plant-Produced HIV-1 Subtype C Envelope gp140 Immunogens. Front Plant Sci, 2019. 10: p. 1378.

20. Maclean, J., et al., Optimization of human papillomavirus type 16 (HPV-16) L1 expression in plants: comparison of the suitability of different HPV-16 L1 gene variants and different cell-compartment localization. J Gen Virol, 2007. 88(Pt 5): p. 1460–9.

21. Sainsbury, F., E.C. Thuenemann, and G.P. Lomonossoff, pEAQ: versatile expression vectors for easy and quick transient expression of heterologous proteins in plants. Plant Biotechnol J, 2009. 7(7): p. 682–93.

22. Tanzer, F.L., et al., The porcine circovirus type 1 capsid gene promoter improves antigen expression and immunogenicity in a HIV-1 plasmid vaccine. Virol J, 2011. 8: p. 51.

